# Repression of pattern-triggered immune responses by hypoxia in Arabidopsis

**DOI:** 10.1101/2023.11.07.565979

**Authors:** Brian C. Mooney, Catherine M. Doorly, Melissa Mantz, Pablo García, Pitter F. Huesgen, Emmanuelle Graciet

## Abstract

Biotic and abiotic stresses frequently co-occur in nature, yet, relatively little is known about how plants co-ordinate the response to combined stresses. Previous research has shown that protein degradation by the ubiquitin/proteasome system is central to the regulation of multiple independent stress response pathways in plants. The Arg/N-degron pathway is a subset of the ubiquitin/proteasome system that targets proteins based on their N-termini and has been specifically implicated in the responses to biotic and abiotic stresses, including hypoxia *via* accumulation of ERF-VII transcription factors, which orchestrate the onset of the hypoxia response program. Here, we investigated the role of the Arg/N-degron pathway in mediating the crosstalk between coinciding abiotic and biotic stresses using hypoxia treatments and the flg22 elicitor of pattern-triggered immunity (PTI), respectively. We uncovered a link between the transcriptional responses of plants to hypoxia and flg22. Combined hypoxia/flg22 treatments showed that hypoxia represses the flg22 transcriptional program, as well as the expression of pattern recognition receptors, MAPK signalling and callose deposition during PTI, through mechanisms that are mostly independent from the ERF-VIIs. These findings aid understanding of the trade-offs between plant responses to combined abiotic/biotic stresses in the context of our efforts to increase crop resilience to global climate change. Our results also show that the well-known repressive effect of hypoxia on innate immunity in animals also applies to plants.

**Significance statement:** Understanding how plants regulate the crosstalk between stress response pathways is key to our efforts to increase crop resilience and mitigate yield losses caused by global climate change. Despite the urgency to do so, relatively little is known about how plants respond to combined stresses, which frequently occur in nature. Here, we show that the hypoxia response program and the basal layer of plant immunity (pattern-triggered immunity or PTI) share components. Our data also show that hypoxia represses several key aspects of PTI, a situation akin to that discovered in animals decades ago. These findings have implications for our ability to develop resilient crops by limiting the negative trade-offs that exist between hypoxia response and immunity.

## Introduction

A key question in biology is ‘how do organisms perceive and respond to environmental cues, as well as stresses?’. While much has been learned about plant responses to individual cues and stresses, the question of how plants integrate information from multiple, combined, stresses to trigger a coordinated response has come to the forefront due to the negative impact of global climate change on crop yields. Studies have shown that the transcriptional response programs to combined stresses can differ considerably from those of the respective individual stresses (Rasmussen et al., 2013; Suzuki et al., 2014; Tan et al., 2023). However, relatively little is known about the mechanisms underpinning such differential transcriptional outputs. The ubiquitin/proteasome system is a key player in coordinating the crosstalk between stress response pathways (reviewed in (Sadanandom et al., 2012; Stone, 2014; Miricescu et al., 2018)) by regulating, for example, the stability of transcriptional regulators that coordinate the crosstalk between phytohormone signalling pathways, such as ethylene/jasmonic acid or jasmonic acid/salicylic acid signalling (Pauwels et al., 2015; Nagels Durand et al., 2016; Miricescu et al., 2018; Wang et al., 2022). The N-degron pathways are a subset of the ubiquitin/proteasome system that target substrate proteins for degradation based on their N-termini (reviewed in (Dissmeyer, 2019; Varshavsky, 2019)). These pathways have been specifically implicated in the responses to a wide range of biotic and abiotic stresses (de Marchi et al., 2016; Gravot et al., 2016; Vicente et al., 2017; Miricescu et al., 2018; Till et al., 2019; Vicente et al., 2019). However, their roles (and that of their substrates) in the regulation of the crosstalk between abiotic/biotic stress response pathways have not been examined.

The arginylation-dependent Arg/N-degron pathway requires the sequential activity of several enzymatic components to modify the N-terminus of substrate proteins before degradation (Fig. 1A). As some of these components are oxygen-dependent, the pathway functions as a *de facto* oxygen sensor that suppresses hypoxia responses in oxygenated conditions *via* the turnover of group VII ETHYLENE RESPONSE FACTOR (ERF-VII) transcription factors, which act as master regulators of the hypoxia response program (Mustroph et al., 2009; Gibbs et al., 2011; Licausi et al., 2011; Reynoso et al., 2019). Other transcriptional regulators, such as LITTLE ZIPPER 2 (ZPR2) and VERNALIZATION2 (VRN2), are also targeted for degradation by the Arg/N-degron pathway in an oxygen-dependent manner (Gibbs et al., 2018; Weits et al., 2019; Labandera et al., 2020). In the case of ERF-VIIs (as well as ZPR2 and VRN2), the N-terminal cysteine (Cys) residue of the proteins is oxidized by PLANT CYSTEINE OXIDASE enzymes (PCOs) (Weits et al., 2014; White et al., 2017; White et al., 2020), followed by the conjugation of arginine (Arg) to their N-terminus by so-called Arg-transferases (ATE1 and ATE2 in *Arabidopsis thaliana*) (Graciet et al., 2009; Zubrycka et al., 2023). This N-terminal Arg (a destabilizing residue) is an N-degron that can be recognized by the PROTEOLYSIS6 (PRT6) E3 ubiquitin ligase, resulting in the degradation of the target proteins (Garzon et al., 2007; Dissmeyer, 2019; Holdsworth et al., 2020; Zubrycka et al., 2023). Arabidopsis *ate1 ate2* (noted *a1a2*) and *prt6* mutants accumulate the ERF-VII transcription factors and consequently display a constitutive activation of the hypoxia response program (Gibbs et al., 2011).

**Figure 1:**
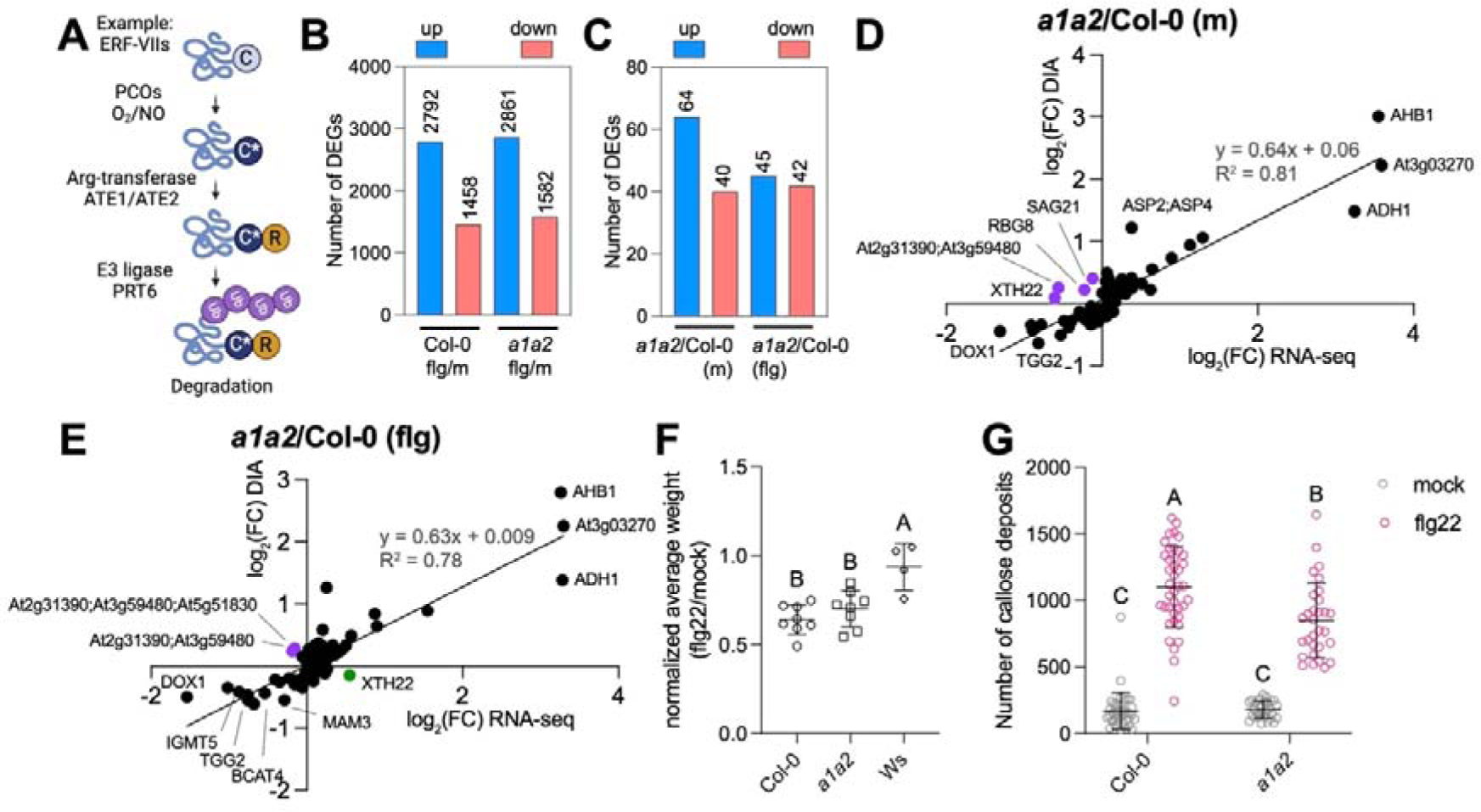
Response of *a1a2* mutant and wild-type Col-0 seedlings to flg22. **(A)** The oxygen-dependent Arg/N-degron pathway. O_2_: oxygen; NO: nitric oxide. **(B)** Number of up and down-regulated DEGs when comparing flg22 to mock-treated samples (flg/m) for wild-type (Col-0) and *a1a2* (a1a2) mutant seedlings. **(C)** Number of up and down-regulated DEGs when comparing *a1a2* to wild-type Col-0 seedlings, either mock (*a1a2*/Col-0 (m)) or flg22-treated (*a1a2*/Col-0 (flg)). **(D)** Comparison of differential protein accumulation and differential gene expression in mock-treated Col-0 and *a1a2* seedlings. **(E)** Comparison of differential protein accumulation and differential expression in flg22-treated Col-0 and *a1a2* seedlings. (D) and (E) Purple: lower mRNA levels and higher protein accumulation in *a1a2* compared to the wild type. Green: higher expression in *a1a2* seedlings, but lower protein levels in this mutant compared to the wild type. The log_2_ of the fold changes (FC) are shown. DIA: Data Independent Acquisition mode. **(F)** Seedling growth inhibition assays. Ws: Wassilewskija (flg22-insensitive control (Zipfel et al., 2004)). Means and standard deviations are shown from 8 independent replicates (8 seedlings/genotype/replicate). Results of statistical tests are shown using the compact letter display (CLD) format after one-way ANOVA with Tukey’s multiple comparison. **(G)** Quantification of callose deposits following flg22 treatment. Means and standard deviations from 4 and 3 biological replicates are shown for Col-0 and *a1a2*, respectively, with 5 seedlings/replicate/condition. The results of two-way ANOVA and Tukey’s multiple comparison tests are shown using CLD.

The Arg/N-degron pathway has also been implicated in the response of plants to biotic stresses, with varying susceptibility/resistance phenotypes (de Marchi et al., 2016; Gravot et al., 2016; Till et al., 2019; Vicente et al., 2019). The *a1a2* and *prt6* mutants were found to be more susceptible to clubroot gall caused by the biotrophic protist pathogen *Plasmodiophora brassicae* and this susceptibility was dependent on the accumulation of the ERF-VIIs in the Arg/N-degron pathway mutant backgrounds (Gravot et al., 2016). Alternatively, overexpression of some ERF-VIIs correlates with increased resistance to the necrotrophic fungus *Botrytis cinerea* (Zhao et al., 2012). This points to a potential double role of ERF-VIIs in the regulation of hypoxia response and plant defenses against pathogens, but their role in plant innate immune pathways or in the crosstalk between hypoxia response and plant immunity have not been explored in detail.

Considering that Arg/N-degron pathway mutants display altered defenses against a range of pathogens with different lifestyles, one hypothesis is that a core aspect of the plant innate immune response, such as pattern-triggered immunity (PTI), is mis-regulated in these mutants. The onset of PTI relies on the detection of conserved Pathogen-Associated Molecular Patterns (PAMPs) at the cell surface by Pattern Recognition Receptors (PRRs) and triggers a transcriptional reprogramming to activate defense-related genes. Here, using flg22 (derived from bacterial flagellin) as a model PAMP and elicitor of PTI, we sought to determine if (i) the Arg/N-degron pathway contributes to the regulation of PTI; (ii) ERF-VIIs play a role in PTI; and (iii) there is a crosstalk between plant responses to hypoxia and to flg22, which could be regulated by the ERF-VIIs. Our transcriptomic analyses of *a1a2* and wild-type seedlings in response to flg22 treatment suggested a transcriptional link between hypoxia response and PTI. The use of combined hypoxia/flg22 treatments further showed that hypoxia represses flg22 transcriptional responses, as well as multiple aspects of PTI, such as MAPK signalling and callose deposition, through mechanisms that are largely independent from the ERF-VIIs. In sum, our data reveal that simultaneous exposure to hypoxia and flg22 has a repressive effect on PTI, which is analogous to the repression of innate immunity by hypoxia in macrophages (Taylor and Colgan, 2017). This finding is also of relevance to the trade-offs between plant responses to abiotic and biotic stresses in our efforts to increase crop resilience to mitigate the yield losses caused by global climate change.

## Results

### Transcriptomic response of *a1a2* seedlings to flg22

To explore whether the Arg/N-degron pathway plays a role in the regulation of PTI, we first used RNA-sequencing (RNA-seq) to compare the genome-wide expression changes of *a1a2* and wild-type seedlings treated with 1 µM flg22 for 1 hr (Fig. S1A). When comparing flg22 to mock-treated seedlings (flg/m), the total number of differentially expressed genes (DEGs; adj. *p*-value<0.05 and |log_2_(FC)|>0.585) was similar in wild-type (Col-0) and in *a1a2* seedlings (∼4,400 DEGs), as well as the number of up- and down-regulated genes (∼2,800 up-regulated and ∼ 1,500 down-regulated genes) (Fig. 1B). Our two datasets had a large overlap with a previously published transcriptomic dataset obtained under similar experimental conditions (Fig. S1B). PTI-associated Gene Ontology (GO) terms were over-represented among the DEGs in wild-type and in *a1a2* seedlings after flg22 treatment (*e.g.* ‘innate immune response’, ‘response to salicylic acid stimulus’, ‘callose deposition in cell wall’, ‘MAPKKK cascade’, or ‘response to oxidative stress’) (Fig. S1C and Suppl. Dataset 1). Differences in the flg22 transcriptional response of the *a1a2* mutant compared to the wild type were also found, with 700 and 507 DEGs being specific to either *a1a2* or the wild type, respectively (Fig. S1B). Neither of these sets of DEGs are enriched for specific GO biological process categories, suggesting that they comprise genes associated with a wide range of functions. We also tested whether there were differences in the amplitude of gene expression changes in DEGs that were common to the two genotypes and found that in response to flg22, the directionality and amplitude of gene expression changes were comparable in *a1a2* and Col-0 (Fig. S1D).

For a more stringent comparison between *a1a2* and wild-type seedlings, we re-analyzed the RNA-seq data by comparing directly the two genotypes under either mock (*a1a2*/Col-0 (m)) or flg22 treatment (*a1a2*/Col-0 (flg)) (cut-offs applied: adj. *p*-value<0.05 and |log_2_(FC)|>0.585). This analysis revealed 104 DEGs in *a1a2*/Col-0 (m) and 97 DEGs for *a1a2*/Col-0 (flg) (Fig. 1C). A GO analysis of all DEGs in *a1a2 versus* wild-type seedlings retrieved GO categories related to known functions of the Arg/N-degron pathway (Fig. S1E and Suppl. Data 2), including ‘response to hypoxia’, ‘regulation of seed germination’, ‘regulation of lipid metabolic process’ and ‘glycoside metabolic process’ (Holman et al., 2009; Gibbs et al., 2011; Gibbs et al., 2014; de Marchi et al., 2016; Zhang et al., 2018). The directionality of the gene expression changes was also as expected, with the expression of several known hypoxia response genes being constitutively induced in mock-treated *a1a2* (Fig. S1F). Of the DEGs identified in both *a1a2*/Col-0 datasets, 59 were differentially expressed in *a1a2* independently of the flg22 treatment (Fig. S1G) and these common DEGs were associated with GO categories known to be constitutively mis-regulated in *a1a2* mutants, such as ‘protein arginylation’ and ‘response to hypoxia’ (Holman et al., 2009; Gibbs et al., 2011; de Marchi et al., 2016). Relatively few genes (28) were specifically mis-regulated in the *a1a2* mutant in response to flg22 (Fig. S1G). For a subset of these genes, we validated that the transcriptional mis-regulation was dependent on the loss of Arg-transferase activity in *a1a2* by using a rescue line (Graciet et al., 2009) with the *ATE1* genomic locus re-introduced to the *a1a2* background (Fig. S1H). Based on the RNA-seq data, the majority (22) of these 28 DEGs had lower expression in *a1a2* compared to the wild type after flg22 treatment (Fig. S1I), with several having known functions in the regulation of plant defenses against pathogens (*e.g. PLANT DEFENSIN2.1* (*PDF2.1*); *VEGETATIVE STORAGE PROTEIN1* (*VSP1*); *CYSTEINE-RICH RLK25* (*CRK25*); *CHITINASE A* (*CHIA*)). Six of these genes were instead expressed at higher levels in *a1a2* compared to the wild type after flg22 treatment.

The *a1a2* mutant is expected to accumulate Arg/N-degron pathway substrates, which could result in protein abundance differences without transcriptional changes. We hence also conducted a proteomic comparison of *a1a2* and wild-type seedlings: 74 proteins accumulated differently in at least one of the four comparisons (*a1a2* (flg/m); Col-0 (flg/m); *a1a2*/Col-0 (m) or *a1a2*/Col-0 (flg); cut-off applied: multi-sample ANOVA with permutation-based FDR< 0.05, followed by Tukeýs *post-hoc* test with FDR<0.05). A comparison of the fold-changes in the proteomic and RNA-seq datasets indicated a positive correlation (Fig. 1D and Fig. 1E). This included proteins/genes up-regulated by hypoxia response (*e.g.* ADH1 and HB1), and proteins involved in glucosinolate biosynthesis (*e.g.* BCAT4, TGG2, IGMT5), which were found to be less abundant in *a1a2* in agreement with their lower expression (de Marchi et al., 2016). A small number of proteins/genes behaved differently in the mock-treated *a1a2* mutant compared to the wild type, with some having known roles in plant defense (*e.g. SENESCENCE-ASSOCIATED GENE 21* (*SAG21*) (Salleh et al., 2012)).

Finally, we examined whether the transcriptional differences identified between flg22-treated *a1a2* and wild-type seedlings could be sufficient to affect PTI. We first tested the effect of prolonged flg22 exposure using seedling growth inhibition (Fig. 1F) and root growth inhibition assays (Fig. S1J). Both indicated that the *a1a2* mutant and the wild type were affected in a similar manner by flg22. The production of apoplastic reactive oxygen species (ROS) upon flg22 treatment was also not affected in the *a1a2* mutant compared to the wild type (Fig. S1K). In contrast, callose deposition assays showed that *a1a2* was negatively affected for callose deposition in response to flg22 treatment (Fig. 1G). This could be a consequence of (i) reduced up-regulation of *GLUCAN SYNTHASE-LIKE5* (*GSL5*), an important callose synthase during flg22 response (Luna et al., 2011), in *a1a2* seedlings (log_2_(flg/m)=0.815) compared to the wild-type (log_2_(flg/m)=1.023); or (ii) *a1a2* having lower levels of glucosinolates (de Marchi et al., 2016), which are important for callose biosynthesis (Clay et al., 2009). Alternatively, the constitutive activation of hypoxia response in *a1a2* could also contribute to the decreased callose deposition phenotype (*e.g.* callose content changes upon hypoxia treatment of wheat seedlings (Subbaiah and Sachs, 2001; Albrecht and Mustroph, 2003)).

### Hypoxia and flg22 transcriptional response programs overlap

Our data suggest a potential connection between hypoxia and flg22 responses. The RNA-seq analysis also revealed that genes associated with the GO categories ‘response to oxygen levels’ and ‘response to hypoxia’ were enriched among flg22-responsive genes in the wild type (Col-0 flg/m) and in *a1a2* (*a1a2* flg/m) (Fig. 2A and Suppl. Dataset 1). We carried out a similar GO analysis using (i) the Arabidopsis flg22 response dataset from Denoux *et al*. (Denoux et al., 2008); and (ii) a dataset composed of conserved flg22-responsive genes across 4 different species of Brassicaceae (termed ‘Brassicaceae core PTI’) (Winkelmuller et al., 2021). A similar enrichment for the ‘response to hypoxia’ and ‘response to oxygen levels’ was found (Fig. 2A), with several hypoxia-related DEGs common to at least two datasets showing the same directionality of expression changes (Fig. S2A). In our Col-0 flg/m dataset, DEGs found in these GO categories included *ERF73/HRE1* (AT1G72360), *ERF71/HRE2* (AT2G47520), *HEMOGLOBIN1* (*HB1*; AT2G16060) and *Alanine AMINOTRANSFERASE1* (*AlaAT1*; AT1G17290), as well as *ENHANCED DISEASE SUSCEPTIBILITY 1* (*EDS1;* AT3G48090) and *PHYTOALEXIN DEFICIENT 4* (*PAD4;* AT3G52430) (Fig. S2B).

**Figure 2:**
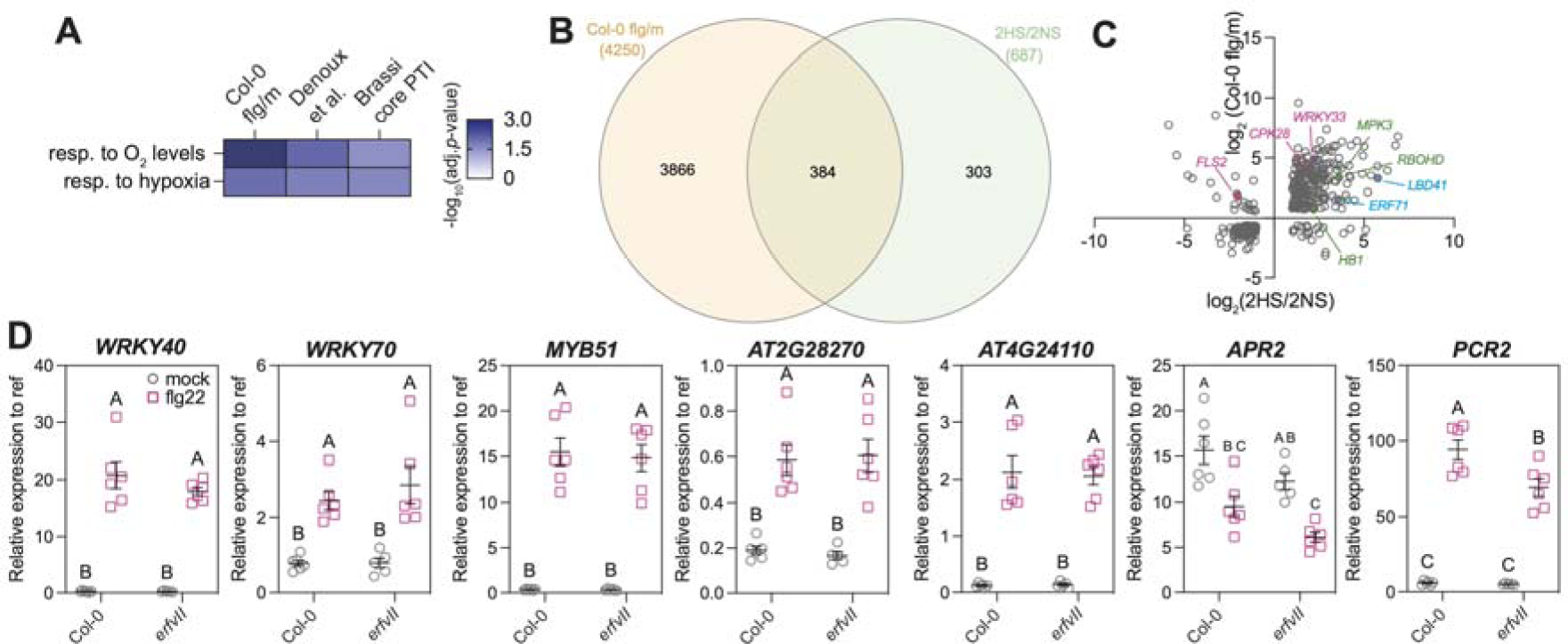
Overlap between the flg22 and hypoxia transcriptional response programs. **(A)** Enrichment for GO categories associated with oxygen levels in different datasets: Col-0 flg/m (this study); Denoux *et al*. (Denoux et al., 2008) and Brassicaceae core PTI (Winkelmuller et al., 2021). The −log_10_ of the corrected *p*-values are shown for the GO categories ‘response to oxygen levels’ (GO#70482) and ‘response to hypoxia’ (GO#1666). See also Suppl. Dataset 1 for a complete list of GO categories and number of DEGs in each GO category. **(B)** Overlap between flg22 response genes identified in Col-0 and differentially regulated genes after 2 hrs of hypoxia (2HS/2NS; O_2_<2%; cut-off applied: |log(FC)|>1.0 and FDR<0.05) (Lee and Bailey-Serres, 2019). **(C)** Comparison of the directionality and amplitude of gene expression change for the 384 genes common to Col-0 flg/m and 2HS/2NS. **(D)** Expression analysis using RT-qPCR of selected genes regulated by both hypoxia and flg22 treatments in wild-type Col-0 and *erfVII* quintuple mutant seedlings mock and flg22-treated. Mean and SEM from 6 biological replicates are shown. The results of two-way ANOVA and Tukey’s multiple comparison tests are shown using CLD.

The overlap between the transcriptional responses to flg22 and to hypoxia was further explored by comparing our Col-0 flg/m dataset with a list of hypoxia responsive genes in 7-day old Arabidopsis seedlings treated with 2 hrs of hypoxia (O_2_ < 2%) compared to normoxia (|log_2_(FC)|>1.0 and FDR<0.05) (Lee and Bailey-Serres, 2019)). This analysis revealed a statistically significant overlap (*p*-value<10^−4^; Chi square test) of 384 DEGs (Fig. 2B and Fig. S3). The majority of these common DEGs had the same directionality of gene expression change in response to hypoxia and to flg22 (Fig. 2C). This was the case for typical flg22 response genes (*e.g. WRKY33* or *CPK28*), hypoxia response genes (*e.g. LBD41* or *HRE2/ERF71*), and genes known to be involved in both response pathways (*e.g. RBOHD, MPK3* or *HB1*). One notable exception was *FLS2*, which codes for the flg22 PRR, and whose transcription was repressed under hypoxia, while being up-regulated upon flg22 treatment. GO analysis of the 384 common DEGs indicated an enrichment for genes associated with defense-related processes (*e.g.* ‘defense response by callose deposition’, ‘indole glucosinolate metabolic process’ or ‘defense response’) and the response to oxygen levels (Suppl. Dataset 4). There was also an over-representation of response genes to other abiotic stresses (*e.g.* osmotic stress, temperature), and to stress-related hormone signalling pathways (*i.e.* ethylene, salicylic acid and jasmonic acid).

To determine if the overlap of hypoxia and flg22 responses could be a conserved feature of the flg22 transcriptional response program, we also compared the Brassicaceae core PTI dataset (Winkelmuller et al., 2021) and hypoxia response genes (Lee and Bailey-Serres, 2019). A statistically significant overlap (*p*-value<10^−4^; Chi square test) of 118 DEGs was found (Fig. S2C), and similarly to the comparison with our Col-0 flg/m dataset, most common genes had the same directionality of gene expression change (Fig. S2D). Hence, the common regulation of a subset of hypoxia and flg22 response genes appears to be a conserved feature, at least within the Brassicaceae family.

Because the ERF-VII transcription factors are the master regulators of the transcriptional response program to hypoxia, we explored whether the common regulation of some genes by both flg22 or hypoxia requires the ERF-VIIs. To this end, the expression of a subset of genes was determined in Col-0 and in an *erfVII* quintuple mutant (Abbas et al., 2015) following mock or flg22 treatment (Fig. 2D). The selected genes were differentially expressed in the 2HS/2NS dataset (Lee and Bailey-Serres, 2019) and in our Col-0 flg/m dataset, and either (i) responded more strongly in *a1a2* flg/m compared Col-0 flg/m; or (ii) were already significantly up-regulated in mock-treated *a1a2* (possibly because of the ERF-VII accumulation). This analysis indicated that the regulation of some hypoxia response genes by flg22 treatment is likely not ERF-VII-dependent.

### Hypoxia represses the flg22 transcriptional response program

To examine the potential crosstalk between hypoxia and flg22 responses, the genome-wide expression changes that occur under individual (hypoxia or flg22) and combined hypoxia/flg22 treatments were determined in wild-type Col-0 seedlings. We focused on a relatively short 1 hr treatment that enabled monitoring of the onset of both flg22 and hypoxia responses (Fig. S4A). The DEGs (cut-offs: adj. *p*-value<0.05 and |log_2_(FC)|>0.585) identified under individual hypoxia (HM v NM; hypoxia/mock v normoxia/mock) or flg22 (NF v NM; normoxia/flg22 v normoxia/mock) treatments compared to untreated seedlings overlapped with those obtained in other similar published datasets (Denoux et al., 2008; Lee and Bailey-Serres, 2019) (Fig. S4B and Fig. S4C), and retrieved an enrichment for expected GO categories (Suppl. Dataset 5). Analysis of the number of DEGs indicated that flg22 treatment alone induced greater transcriptional changes (NF v NM; 1906 DEGs) than hypoxia alone (HM v NM; 145 DEGs), likely because of the short 1 hr treatment (Fig. 3A). Combining hypoxia/flg22 resulted in even larger transcriptional changes (HF v NM; 2612 DEGs) than flg22 treatment alone (Fig. 3A). A comparison of the three datasets showed a large overlap between the responses to flg22 alone (NF v NM) and to combined hypoxia/flg22 treatment (HF v NM; 1601 DEGs in common), and a smaller overlap between hypoxia and combined hypoxia/flg22 treatments (131 DEGs) (Fig. 3B). In addition, 59 genes were commonly differentially expressed in all conditions, with an over-representation of genes associated with metabolic processes (*e.g.* ‘response to carbohydrate stimulus’), hormone signalling (ethylene, abscisic acid and brassinosteroids), immunity (‘response to chitin’) and development (‘root hair elongation’ or ‘developmental maturation’) (Suppl. Dataset 5). Fifty five of these 59 DEGs showed the same directionality of gene expression change irrespective of the stress applied (Fig. S4D). For the 53 genes that were up-regulated in all conditions (Fig. 3C), the combined hypoxia/flg22 treatment enhanced the amplitude of up-regulation (Fig. 3D and Fig. 3E). While there were only 2 common genes being down-regulated (Fig. 3F), the same trend was observed, such that combined hypoxia/flg22 resulted in a stronger repression of these 2 genes compared to hypoxia or flg22 alone (Fig. 3G).

**Figure 3:**
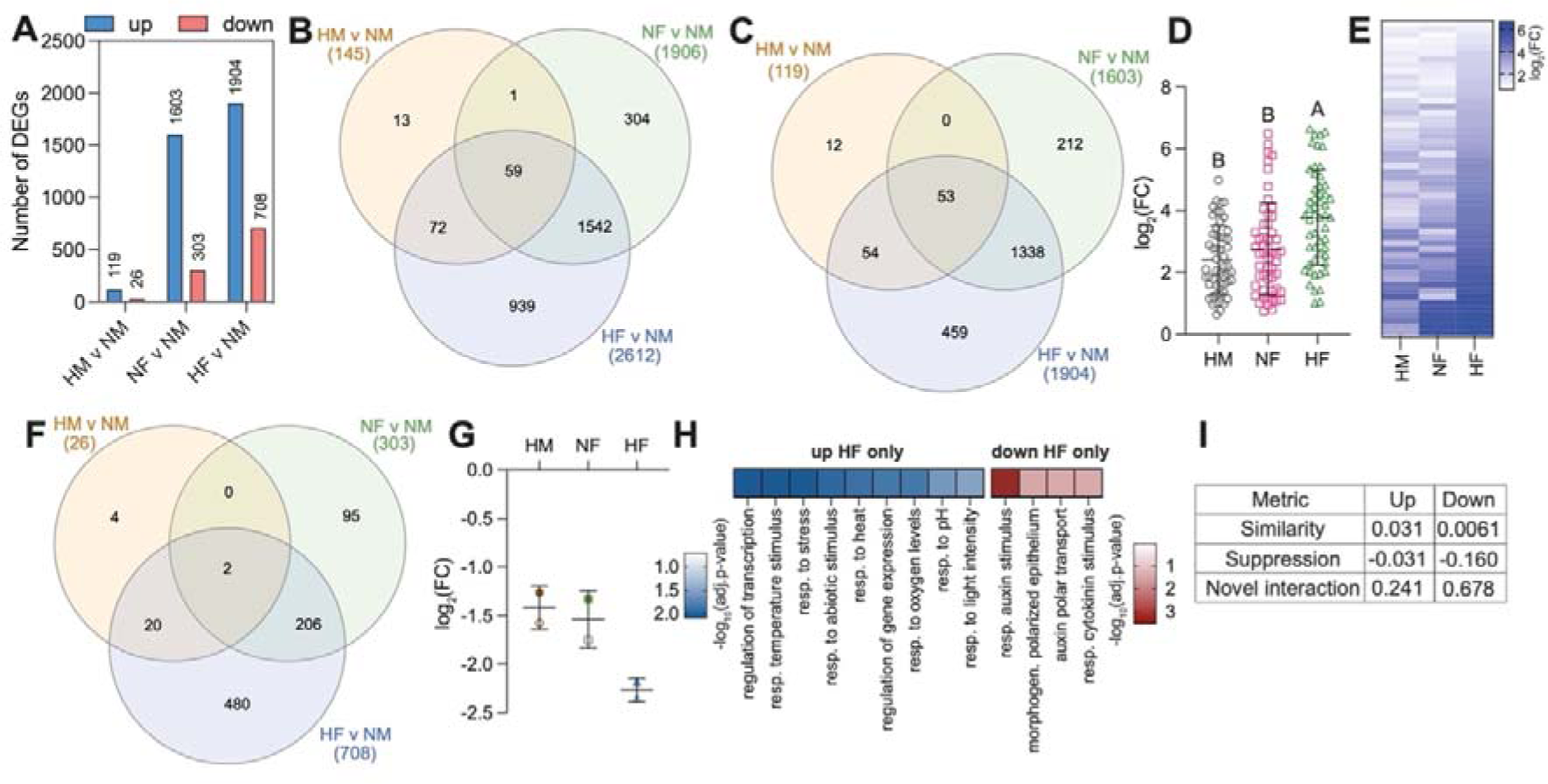
Effects of combined hypoxia/flg22 treatments on genome-wide transcription. **(A)** Number of up- and down-regulated genes. NM: normoxia/mock; HM: hypoxia/mock; NF: normoxia/flg22; HF: hypoxia/flg22. **(B)** Overlap between the 3 different datasets using all DEGs. **(C)** Overlap between the up-regulated DEGs in each of the 3 datasets. **(D)** Average expression of the 53 up-regulated DEGs common to all 3 datasets. Results of statistical tests are shown after one-way ANOVA with Tukey’s multiple comparison. **(E)** Heatmap showing the log_2_(FC) in HM, NF and HF-treated seedlings for the 53 up-regulated DEGs common to all 3 datasets. **(F)** Overlap between the down-regulated DEGs. **(G)** Average expression of the 2 down-regulated DEGs common to all 3 datasets. Each gene is respectively represented with filled or open symbols. **(H)** Selected GO terms enriched (corrected *p*-value<0.05) amongst uniquely up- and down-regulated DEGs upon combined hypoxia/flg22. See Suppl. Dataset 5 for full list of GO terms and number of DEGs in each GO category. **(I)** Metrics to characterize the transcriptional interaction between individual (hypoxia or flg22) stresses and combined hypoxia/flg22 treatment (see Fig. S4 for calculations).

Strikingly, 939 genes were uniquely regulated in response to combined hypoxia/flg22 (HF v NM), indicating that the combined stress elicits a distinct transcriptional response. GO analysis of these 939 hypoxia/flg22-specific DEGs revealed an enrichment for genes associated with expected defense and phosphorylation-related functional GO categories. GO analysis for molecular functions revealed an over-representation of transcription factor-coding genes (Suppl. Dataset 5), with bHLH, Zn finger and homeobox transcription factors accounting for a larger number of differentially expressed transcription factors (Fig. S4E). The expression of these transcription factors was mostly repressed by hypoxia/flg22 treatment, while NAC, WRKY and bZIP transcription factors were up-regulated. In contrast, the 304 DEGs specific to flg22 treatment did not have an over-representation of transcription factor coding genes, so this appears to be a unique feature of the response program to combined hypoxia/flg22 (Suppl. Dataset 5). Separate GO analyses for uniquely down-regulated genes upon hypoxia/flg22 revealed an enrichment for terms associated with developmental processes, such as ‘morphogenesis of polarized epithelium’, including genes associated with phytohormone signalling pathways with roles in cell division and differentiation (*i.e.* auxin and cytokinin signalling) (Fig. 3H). In contrast, genes that were uniquely up-regulated after the combined treatment were enriched for genes associated with stress responses (*e.g.* response to heat), suggesting that the response to combined stresses could be broader and overlap with that of other individual stresses (Fig. 3H).

To further characterize the crosstalk between hypoxia and flg22 responses, we used three metrics established to study the crosstalk between the transcriptional programs to combined abiotic stresses in *Marchantia polymorpha* (Tan et al., 2023) (see also Fig. S4F for calculations). Specifically, these metrics allow the quantification of (i) shared aspects of the transcriptional response programs (similarity score ranges from 0 to 1, with 1 corresponding to identical response programs), (ii) stress dominance (suppression score varies between −1 and 1, with a negative score indicating a suppression of flg22 response by hypoxia), and (iii) novel components of combined treatments (novel interaction score; varies between 0 and 1, with 1 indicating that combined stresses results in an entirely new transcriptional program) (Tan et al., 2023). For both up- and down-regulated genes, the negative suppression score obtained suggests that the hypoxia response program could repress flg22 transcriptional response (Fig. 3I). The lower suppression score obtained for down-regulated genes (−0.16) suggests that the repression of down-regulated genes by hypoxia could be of particular importance. The novel interaction scores obtained further indicate that combined hypoxia/flg22 elicits a transcriptional response that is different from each of the individual stresses. The novel interaction score for up-regulated genes (0.241) is in line with the average novel interaction scores for up-regulated genes in response to 18 combined treatments in Marchantia (0.29). Notably, the novel interaction score for down-regulated genes (0.678) is higher than in Marchantia, in which the highest score obtained for down-regulated genes was 0.56 (18 treatments). This indicates that the repression of specific genes under combined hypoxia/flg22 might be a particularly important aspect of the unique response to this combined stress. A GO analysis of the 480 down-regulated genes under hypoxia/flg22 only (Fig. 3H) suggests an over-representation for genes associated with development. Strikingly, there is also an enrichment for genes coding for cytochrome P450 enzymes, which use molecular oxygen to catalyze the oxidation of other molecules (Suppl. Dataset 5).

The metrics derived from our RNA-seq dataset analysis suggest that hypoxia represses aspects of flg22 response. We therefore examined more carefully the 304 DEGs (Fig. 3B) identified only after flg22 treatment (*i.e*. these genes which no longer respond to flg22, when combined with hypoxia). A GO analysis of these DEGs showed an enrichment for molecular functions associated with phosphorylation (‘kinase activity’), ‘carbohydrate binding’ and ‘transmembrane receptor protein kinase’. Notably, genes in the latter GO category included PRRs such as *FLS2* and *ELONGATION FACTOR Tu RECEPTOR* (*EFR*). Hence, our data suggest that one particular aspect of the repression of flg22 response by hypoxia could occur at the level of PRR expression.

### Hypoxia suppresses PTI

Next, we tested whether different aspects of PTI were also repressed by combined hypoxia/flg22, and examined if ERF-VII transcription factors played a role in the repression of PTI under hypoxic conditions. We first used root growth inhibition assays to reveal additive or synergistic effects of the combined treatment (Fig. 4A) because either flg22 or hypoxia suppress root elongation. Either flg22 (NF) or hypoxia (HM) treatments resulted in decreased root elongation compared to the same genotype in control conditions (normoxia/mock (NM)), with hypoxia having a stronger inhibitory effect than flg22. In Col-0, the combined hypoxia/flg22 (HF) treatment resulted in roots that were shorter than those of seedlings treated with flg22 alone. Compared to hypoxia alone (HM), combined hypoxia/flg22 treatment did not impair further root elongation of Col-0 seedlings. In contrast, the *erfVII* quintuple mutant was more severely affected by the combined HF stress compared to both HM and NF conditions, thus suggesting that ERF-VIIs in the wild type may contribute to protecting the root meristem under combined hypoxia/flg22 treatment. Expression analysis by RT-qPCR of selected immunity-related genes confirmed that hypoxia dampens the up-regulation of flg22 response genes (Fig. 4B), including PRR-coding genes such as *FLS2* and *EFR*, through mechanisms that appear to be mostly independent from the ERF-VIIs. Immunoblot analysis of FLS2 protein levels using seedlings of a *FLS2_pro_:FLS2-3xmyc-GFP* line (Robatzek et al., 2006) further suggested that FLS2 protein levels are unchanged after 1 hr of combined hypoxia/flg22 compared to flg22 treatment (Fig. S5A). As expected, in Col-0, phosphorylation of MPK3 and MPK6 was induced after 30 min of flg22 treatment under normoxia (NF). We did not observe a detectable accumulation of phosphorylated MPKs by immunoblot after a 30-min hypoxia treatment alone, likely because MPK3/6 activity peaks at 2 hrs of hypoxia and is only weakly activated at 30 min, as shown using more sensitive radioactive in-gel kinase assays (Chang et al., 2012). Notably, the level of phosphorylated MPK3/6 was reduced under combined hypoxia/flg22 treatment compared to flg22 treatment alone. Similar effects were observed in the *erfVII* mutant (Fig. 4C, Fig. 4D and Fig. S5B). In contrast, *MPK3/4/6* expression was not repressed under hypoxia/flg22 compared to flg22 treatment alone (Fig. S5C). Finally, callose deposition was significantly decreased under combined hypoxia/flg22 conditions in Col-0 and *erfVII* seedlings, compared to treatment with flg22 alone (NF) (Fig. 4E and Fig. S5D), with no statistically significant differences between Col-0 and *erfVII.* Altogether, our data indicate that several aspects of PTI are repressed under simultaneous hypoxia/flg22 treatments compared to flg22 alone, and that these effects are mostly ERF-VII independent.

**Figure 4:**
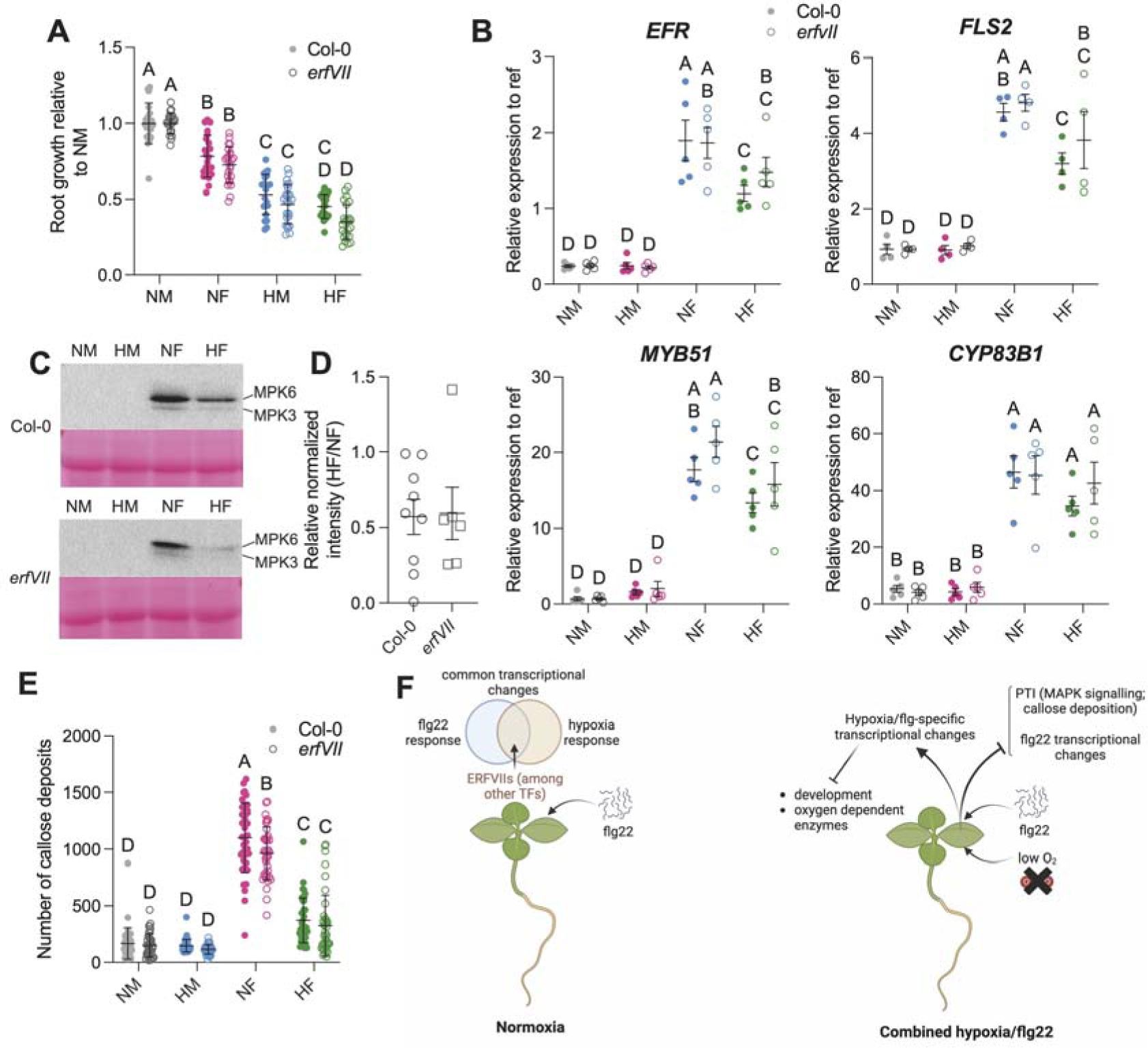
Repression of PTI by hypoxia in combined hypoxia/flg22 experiments. **(A)** Root growth inhibition assays with wild-type (Col-0) and *erfVII* mutant seedlings under combined hypoxia/flg22 treatment. Means of the normalized root growth relative to untreated conditions (NM) and standard deviations of 4 biological replicates with 5-6 seedlings/genotype/condition in each biological replicate are shown. The results of statistical tests (two-way ANOVA with Tukey’s test) are shown. **(B)** Transcriptional suppression of immunity-related genes under combined hypoxia/flg22 treatment compared to normoxia/flg22 in Col-0 and *erfVII*. Means and SEM from 5 biological replicates are shown. The results of two-way ANOVA and Fisher’s test are shown. **(C)** Anti-phosphorylated MPK immunoblots using Col-0 and *erfVII* mutant seedlings. See also Fig. S5. **(D)** Relative intensity of MPK signals under combined hypoxia/flg22 compared to normoxia/flg22 after normalization with Ponceau intensity. Mean and SEM of 9 (Col-0) and 6 (*erfVII*) biological replicates are shown. **(E)** Quantification from callose deposition experiments. Mean and standard deviation from 4 biological replicates are shown, with 5 seedlings/genotype/condition in each biological replicate. The results of two-way ANOVA and Tukey’s tests are shown. Note that the Col-0 NM and Col-0 NF data is also shown in Fig. 1G. **(F)** Model summarizing the crosstalk between hypoxia and PTI under normoxia, and upon combined hypoxia/flg22.

## Discussion

In animals, the complex crosstalk between hypoxia and immunity has been known for decades and some of the underlying mechanisms have been dissected in different cell types. For example, using murine macrophages as a model system, hypoxia has been shown to dampen innate immune signalling pathways (Rahat et al., 2011; Taylor and Colgan, 2017; Weigert et al., 2018). In plants, it has been shown that infection with the necrotrophic fungus *Botrytis cinerea* can result in localized hypoxic niches as a consequence of increased respiration and/or water soaking (Valeri et al., 2021). Other pathogens trigger local hypoxic environments *via* the formation of galls/tumors (e.g. *Agrobacterium tumefaciens*) (Kerpen et al., 2019). In this context, the ERF-VII transcription factors have been shown to play a role in the ability of plants to defend themselves (Gravot et al., 2016; Kerpen et al., 2019). Despite these potential links between hypoxia and infection by necrotrophic and gall-forming pathogens, the connection between hypoxia and innate immune signalling pathways has not been investigated in detail. Here, we used transcriptomics datasets to identify connections between hypoxia and PTI, and show that hypoxia represses PTI in Arabidopsis. We also examined whether ERF-VIIs play a role in the crosstalk between hypoxia response and PTI.

The genome-wide comparison of the transcriptional changes induced by flg22 treatment in wild-type Col-0 and *a1a2* seedlings revealed that the flg22 transcriptional response program includes sets of hypoxia response genes, and that this overlap is a conserved feature, at least within the Brassicaceae (see model in Fig. 4F). Our transcriptomic data with the *a1a2* mutant, that constitutively accumulates ERF-VII transcription factors (Gibbs et al., 2011) suggested that ERF-VIIs could play a role in the transcriptional regulation of some flg22-responsive genes. Despite the fact that a proportion of these flg22-responsive genes were bound by HRE2 (Lee and Bailey-Serres, 2019), our RT-qPCR data instead suggests that ERF-VIIs are unlikely to play a role.

A key question is whether the hypoxia-related genes identified in flg22 transcriptomics datasets are regulated due to the formation of a hypoxic environment during PTI, or require alternative hypoxia-independent mechanisms for their regulation. Our data indirectly suggest that flg22 treatment does not trigger hypoxia in seedlings, as we did not detect the up-regulation of many well-characterized hypoxia marker genes, such as *ALCOHOL DEHYDROGENASE1* (*ADH1*) or *PLANT CYSTEINE OXIDASE1* (*PCO1*). This is in agreement with previous data suggesting that flg22 concentrations of up to 10 µM (a concentration higher than those used in this study) are not sufficient to trigger local hypoxic niches in 3-4-week old plants (Valeri et al., 2021). Hence, the regulation of common genes by either hypoxia or flg22 treatment may not require the formation of a hypoxic environment, but instead might involve other mechanisms.

Considering the overlap between the transcriptional responses to flg22 and hypoxia, and also the fact that hypoxic niches form during infection by necrotrophic and gall-forming pathogens (Kerpen et al., 2019; Valeri et al., 2021), we examined more carefully the potential crosstalk between hypoxia and PTI by first comparing the genome-wide transcriptional changes in wild-type seedlings treated with individual stresses (hypoxia or flg22 alone) or combined hypoxia/flg22. The use of metrics developed to compare the crosstalk between stresses (Tan et al., 2023) revealed that hypoxia could repress the transcriptional flg22-response program. For example, 304 differentially expressed genes upon flg22 treatment were no longer transcriptionally regulated when seedlings were subjected to combined hypoxia/flg22 (Fig. 3B).This repression is particularly relevant for genes that are down-regulated by flg22. Indeed, the negative suppression score was lower for down-regulated genes (−0.16 compared to −0.031 for up-regulated genes), and importantly was also within the same range as the suppression scores obtained when combined abiotic stresses were applied to *Marchantia* (Tan et al., 2023). One possible explanation is that plants may experience resource limitations that result in the prioritization of their transcriptional response to hypoxia compared to the onset of PTI, perhaps because hypoxia could have a more immediate impact on plant survival.

Unique features associated with the response of wild-type seedlings to combined hypoxia/flg22 emerged in our datasets, with over 900 unique DEGs upon hypoxia/flg22 treatment, compared to hypoxia or flg22 treatments alone. The novel interaction score obtained for down-regulated genes upon combined hypoxia/flg22 was higher than the novel interaction scores found for the range of 18 combined abiotic stresses in Marchantia (Tan et al., 2023), thus suggesting that the repression of specific sets of genes is an important and unique feature of plant responses to combined hypoxia/flg22. These unique features include the repression of developmental processes, which could correlate with a resource/allocation problem and potential energy crisis that is typical of hypoxia stress. In addition, we observed the repression of genes coding for enzymes (cytochromes P450) that use oxygen as co-substrate, suggesting a limitation of enzymatic processes that consume oxygen (Fig. 4F).

Another unique feature of the combined hypoxia/flg22 response was the over-representation of genes associated with transcription factors, with some families being up-regulated (NAC, WRKY, bZIP), while others tended to be predominantly down-regulated (bHLH, Zn finger, homeobox). Altogether, the data suggest that the coordination of plant responses to combined hypoxia/flg22 requires the orchestration of gene expression changes under the control of specific transcription factor families. Notably, some of the up-regulated transcription factor families, such as NAC or WRKY, are typically associated with stress responses (Shao et al., 2015). The WRKY22 transcription factor had previously been identified as being up-regulated during submergence in the dark (which results in hypoxia) and to play a role in the induction of plant defenses to pathogens following a 12-hr dark submergence treatment (Hsu et al., 2013). These stress conditions are different from our combined hypoxia/flg22 treatments. Nevertheless, we found *WRKY22* up-regulated in our hypoxia/flg22 dataset, together with 11 (out of 45) genes whose genomic regions were shown to be bound by WRKY22 (Hsu et al., 2013). This indicates that WRKY22 may be a regulator of plant responses to combined hypoxia/flg22.

Some of the transcription factor families that are mostly repressed (*e.g.* homeobox transcription factors) are generally associated with developmental processes (Jeong et al., 2012). This suggests again an increased need to regulate resource allocation under combined stress, with a stronger prioritization of stress response pathways as opposed to developmental processes. Our core findings with combined hypoxia/flg22 (*i.e.* the preponderance of a novel transcriptional response, the relevance of transcription factors and the stronger prioritization of stress responses) are reminiscent of previous findings in studies with combined abiotic/biotic stresses (Atkinson and Urwin, 2012; Prasch and Sonnewald, 2013; Rasmussen et al., 2013; Suzuki et al., 2014; Pandey et al., 2015; Zhang and Sonnewald, 2017; Saijo and Loo, 2020). However, such studies have thus far focused on combinations between pathogens and abiotic stresses such as heat and drought, and have not included hypoxia.

Notably, the repression of PTI by hypoxia is not observed just at the transcriptional level. Our data show that MAPK signalling and callose deposition are repressed under combined hypoxia/flg22. In the case of MAPK, the inhibitory effects appear to be independent of the transcriptional regulation of *MPK3/4/6*, and occurs either post-transcriptionally on MPK proteins, or further upstream in the flg22-dependent signalling events originating from FLS2. Similarly, the expression of genes encoding key PTI components, such as the PRRs *FLS2* and *EFR*, is repressed under combined hypoxia/flg22 compared to flg22 alone. However, RT-qPCR analysis using *erfVII* mutants suggested that these master regulators of the hypoxia response program do not play a major role in regulating plant responses to combined hypoxia/flg22. Potential other candidates include WRKY22 (Hsu et al., 2013) as discussed above, or the ZPR2 and VRN2 transcriptional regulators, which are also known to be degraded by the N-degron pathway in an oxygen-dependent manner (Gibbs et al., 2018; Weits et al., 2019; Labandera et al., 2020). VRN2 degradation also depends on the presence of nitric oxide (Gibbs et al., 2018), an important signalling molecule induced in response to both hypoxia and PAMPs/pathogens (Mugnai et al., 2012; Bleau and Spoel, 2021; Borrowman et al., 2023). The role of nitric oxide in integrating hypoxia and immune signaling thus appears as an interesting focus for further studies.

Connections and crosstalk between hypoxia response and immunity/plant defenses are also likely to evolve in the course of development, as the levels of ERF-VIIs and the regulation of innate immunity change with developmental stages and age (Kus et al., 2002; Lozano-Duran and Zipfel, 2015; Giuntoli et al., 2017; Giuntoli and Perata, 2018; Zou et al., 2018). Such developmentally-dependent crosstalk between abiotic and biotic stresses was identified in the context of combined mild salt stress and pathogen infection with the biotrophic oomycete *Hyaloperonospora arabidopsidis* or with the hemibiotrophic bacterium *Pseudomonas syringae* (Berens et al., 2019). In the latter study, the hormonal crosstalk between abscisic acid (ABA) and salicylic acid (SA) played an important age-dependent role. The hypoxia/immunity crosstalk may also be particularly complex because, in contrast to other abiotic stresses, hypoxia itself has a dual nature, in that it may be considered as either a stress (*e.g.* during flooding; termed ‘acute hypoxia’ in this context), or a physiological condition (*e.g.* in meristems, which are hypoxic; termed ‘chronic hypoxia’) (Loreti and Perata, 2020; Weits et al., 2021). Hence, the output of the crosstalk between hypoxia and immunity could differ between tissues and cell types, as observed in mammals (reviewed in (Taylor and Colgan, 2017)) and as suggested by a recent single-cell RNA-seq study (Liu et al., 2023).

Finally, combined hypoxia/flg22 treatment indicates that plants or tissues experiencing acute hypoxia are affected for PTI, which could be relevant in terms of agricultural applications, as it could suggest that flooding may have a negative impact on the ability of plants to fight-off pathogens. This could be exacerbated by the fact that flooding also causes an increased risk of pathogen infection, likely as a result of increased dampness and changes to the soil and plants’ microbiome (Hartman and Tringe, 2019; Gschwend et al., 2020). Hence, the crosstalk between hypoxia and immunity is a new trait to consider in our endeavour to generate more climate resilient crops.

## Materials and Methods

### Arabidopsis lines used

*Arabidopsis thaliana* accession Columbia-0 (Col-0) was used for this study. Mutant lines used are listed and described in Suppl. Table S1. *Arabidopsis* accession Wassilewskija (Ws) was used as a flg22-insensitive control (Zipfel et al., 2004).

### Plant growth conditions

*A. thaliana* plants were grown in 4-cell pots on a soil mixture containing a 5:3:2 ratio of compost:vermiculite:perlite. The soil mixture was sterilized by autoclaving prior to use. When axenic conditions were required, seedlings were grown in Petri dishes containing 0.5xMS medium (pH 5.7) and 6 g/L agar with 0.5% sucrose (w/v). Prior to sowing on agar medium, seeds were surface-sterilized using the vapor-phase sterilization method (Lindsey et al., 2017). Trays or plates were kept at 4°C for 3 days prior to transfer to growth rooms.

### *Arabidopsis thaliana* flg22 RNA-Seq experiments

Seedlings were grown on 0.5xMS agar with 0.5% sucrose plates in continuous light at 20°C. After 9 days of growth, 50 seedlings *per* genotype *per* treatment were transferred to 2 wells of a 6-well plate, each containing 6 mL of 0.5xMS with 0.5% sucrose liquid medium and returned to continuous light with gentle agitation at 120 rpm. 24 hrs after transfer, 1 µM flg22 or an equivalent volume of water (mock treatment) were added to each well. After 1 hr, all seedlings were harvested in liquid nitrogen and stored at −80°C. After grinding, frozen tissue was divided equally for RNA extraction (for RNA-seq) and for protein isolation (proteomic analysis).

### RNA-seq for combined hypoxia/flg22 experiments

Seedlings were grown on 0.5xMS agar supplemented with 0.5% sucrose in continuous light at 20°C for 9 days. On day 9, five seedlings *per* genotype *per* treatment were transferred to 35 mm x 10 mm dishes, each containing 4 mL of 0.5xMS liquid medium and placed on a shaker at 120 rpm overnight in continuous light conditions (20°C). On day 10, seedlings were treated simultaneously with 100 nM flg22 or equivalent volume of water (mock), and with hypoxia (in anaerojars) or normoxia at ambient O_2_ levels. After 1 hr, the seedlings were harvested in liquid nitrogen and stored at −80°C.

### Protein extraction and proteomics sample preparation

Protein extraction buffer (2% SDS, 100 mM HEPES pH 7.5, 5 mM EDTA, Thermo Halt Cocktail 1:100) was added to the ground tissue, which originated from the same samples used for RNA-seq (i.e. the powder was split in 2 for both RNA-seq and proteomics). The volume of buffer added was 1 µL for 0.3 mg of tissue, with a minimum volume of 300 µL *per* sample. Proteins were then denatured by heating samples at 70°C for 12 minutes and samples were centrifuged at 2,500 x g for 10 minutes. The protein lysate was transferred to a fresh tube and protein was quantified using the amido black protein quantification method (Schaffner and Weissmann, 1973; Popov et al., 1975).

For proteome analysis, 20 µg each sample were treated with 0.5 µl benzonase (∼26 U/µl) at 37°C for 30 min, reduced with 10LmM dithiothreitol (DTT) shaking at 37°C for 30 min and alkylated with 50LmM chloroacetamide (CAA) at room temperature in the dark for 30 min. The reaction was quenched with an additional 50LmM DTT for 20 min before protein purification with SP3-beads (Cytiva) as described previously (Hughes et al., 2019). The samples were resuspended in 100 mM HEPES, 2.5 mM CaCl_2_, pH 7.5 and digested with trypsin (proteome:enzyme ratio 100:1 w/w) overnight at 37°C. Digested peptides were desalted with self-packed C18 STAGE-tips (Rappsilber et al., 2007) and resuspended in 0.1% formic acid (FA).

### Mass spectrometry and proteomics data analysis

LC–MS/MS analysis was performed with an UltiMate 3000 RSCL nano-HPLC system (Thermo) coupled to an Impact II Q-TOF mass spectrometer (Bruker) with a CaptiveSpray ion source (Bruker) operated with an acetonitrile (ACN)-saturated nitrogen gas stream. For each run, an estimated 1 μg peptides were loaded on an μPAC pillar array trap column (1 cm length, PharmaFluidics) and separated on a μPAC pillar array analytical column (50 cm flowpath, PharmaFluidics). Peptides were eluted using a 2Lhr elution protocol that included an 80 min separation gradient from 5% to 32.5% solvent B (solvent A: 0.1% FA, solvent B: 0.1% FA in ACN) at a flow rate of 600LnlLmin^−1^, with columns temperated at 40°C. Tandem mass spectra were acquired at 15 Hz in Data Independent Acquisition (DIA) mode, with 400–1200 precursor m/z windows for MS1 and 32 windows of 25Lm/z with 0.5Lm/z overlap in a range from 200–1750 m/z for MS2.

Peptide sequences were identified with DIA-NN (Demichev et al., 2020), version 1.8, using a spectral library predicted from the UniProt *Arabidopsis thaliana* reference proteome (UP000006548_3702, release 2021_02). Search settings were set to double-pass mode with a precursor FDR of 1%, considering Cys carbamidomethylation as a fixed modification. Downstream data analysis was performed using the Perseus software (Tyanova et al., 2016), v.1.6.15.0. Data was log2-transformed and grouped by treatments and genotype in 4 groups (*ate1 ate2* or wild type treated with flg22 or control). Multi-sample ANOVA (permutation-based FDR<0.05) followed by Tukey’s *post-hoc* test with FDR<0.05) as implemented in Perseus was used to determine significant difference in abundance between the conditions.

### Flg22 seedling growth and root growth inhibition assays

Seedlings were grown on 0.5xMS agar plates supplemented with 0.5% sucrose in continuous light at 20°C.

For seedling growth inhibition assays: individual 5.5-day old seedlings were transferred to one well of a 48-well plate containing 1 mL of liquid 0.5xMS medium supplemented with 100 nM flg22 (or equivalent volume of water (mock). For each condition and genotype, 8 seedlings were used in a given biological replicates. Seedlings were then grown with mild shaking (120 rpm) in continuous light (20°C) for 7 days. To weigh the seedlings, all 8 seedlings for a given genotype and condition were harvested together on a paper towel to remove any liquid and weighed together.

For root growth inhibition assays to compare Col-0 and *ate1 ate2*: 5.5-day old seedlings were transferred to 0.5xMS with 1% agar plates that contained either a mock solution (water) or 500 nM flg22 and grown vertically for 3 days in continuous light (20°C). Root elongation during this 3-day period was measured using Image J.

For root growth inhibition assays under combined hypoxia/flg22: 5.5-day old seedlings were transferred to 0.5xMS with 1% agar plates that contained either a mock solution (water) or 500 nM flg22, followed by vertical growth in short-day conditions (8 hrs light; 16 hrs dark) for 48 hrs under normal oxygen conditions. Hypoxia treated seedlings were kept at 0.5% oxygen (applied using PhO_2_X Box (Baker Ruskinn)); plates kept vertical) for 48 hrs in short-day conditions. Normoxia-treated seedlings were kept under normoxic and short-day conditions for 48 hrs. After this 48-hr hypoxia/normoxia treatment, all seedlings were kept vertical for an additional 3 days in short-day conditions and normoxia (*i.e.* recovery period). Root elongation during the 3-day recovery period was determined using ImageJ.

### Reverse transcription quantitative PCR (RT-qPCR)

RNA was reverse transcribed using RevertAid Reverse Transcriptase (Thermo Fisher) in the presence of RiboLock RNase inhibitor (Thermo Fisher), oligo(dT)18 and 1 mM dNTP mixture at 42°C for 45 minutes. qPCR reactions were carried out in a LightCycler 480 instrument (Roche) using 1 µL of cDNA mixed with 1 µL of a primer pair mixture (1 μM final concentration each), 5 µL 2X SYBR green master mix (Roche) and nuclease-free water added to a final volume of 10 μL *per* well. The second derivative maximum method was used to determine crossing point (Cp) values. Gene expression was calculated relative to a reference gene with the comparative Ct method (Cp_reference gene_ – Cp_gene of interest_ = deltaCp). Assuming a PCR efficiency value of 2, relative expression was calculated as 2^deltaCp^. *MON1* (AT2G28390) was used as a reference gene for RT-qPCRs (de Marchi et al., 2016). Primers used for RT-qPCR analysis are listed in Suppl. Table S2.

### RNA-Sequencing, data processing and functional analysis

Each RNA-seq experiment was conducted using samples from three biological replicates. Total RNA was isolated using the Spectrum Total RNA kit (Merck), and RNA integrity was assessed using an Agilent 2100 Bioanalyzer (Agilent). All RNA samples had RNA integrity (RIN) values >7.0. Library preparation and paired-ended 100 bp next-generation sequencing was performed by BGI (Hong Kong) using the BGI-seq PE100 platform. Data processing was carried out by BGI using the filtering software SOAPnuke (including removal of reads containing the adaptor; removal of reads whose N content is greater than 5%; and removal of low-quality reads). The Hierarchical Indexing for Spliced Alignment of Transcripts (HISAT) software was then used for mapping clean reads to the reference genome. Differential gene expression was determined using DESeq2. Cut-offs of adj. *p*-value<0.05 and |log2(FC)|>0.585 were applied to determine differentially expressed genes (DEGs). Significant enrichment analysis of GO function on DEGs was carried out using the BiNGO plug-in in Cytoscape. Overlap between datasets was determined using InteractiVenn (Heberle et al., 2015), and statistical significance of the overlap between datasets was calculated using 2×2 contingency tables and Chi-square tests.

### Callose deposition assays

Seedlings were grown on 0.5xMS agar with 0.5% sucrose plates for 9 days under short-day light conditions (8 hrs light; 16 hrs dark) at 20°C. After 9 days of growth, 5 seedlings *per* genotype *per* treatment were transferred to 3 mL 0.5xMS liquid medium in 35 mm x 10 mm petri dishes. These were placed on a shaker at 120 rpm in short-day conditions before the dark period. On day 10, seedlings were treated for 24 hrs as follows: NM = normoxia (ambient O_2_) and mock treatment (addition of a volume of PBS equivalent to that of the flg22 solution); NF = normoxia (ambient O_2_) and flg22 (1 µM; prepared in PBS); HM = hypoxia (5% O_2_) and mock treatement; HF = hypoxia (5% O_2_; applied using PhO_2_X Box (Baker Ruskinn)) and 1 µM flg22. After 24 hrs, treatment solutions were aspirated, 100% ethanol was added and samples were returned to short days on a shaker at 120 rpm. After 24 hrs, the ethanol was removed and seedlings were washed with 0.07 M PBS (pH 9). Seedlings were stained using 0.01% (w/v) aniline blue in 0.07 M potassium phosphate buffer (pH 9) for 2 hrs with constant agitation at 120 rpm. Cotyledons were mounted on slides using 50% (v/v) glycerol and imaged using fluorescent microscope under DAPI filter. Quantification of callose deposits was done as detailed in (Mason et al., 2020). Briefly, the Trainable Weka Segmentation (TWS) plugin on Fiji was used to identify callose deposits while excluding background fluorescence (*e.g.* vasculature or edges of tissue), by manually training the plugin using a good example of callose deposition. Once trained, all pictures obtained were analyzed using the plugin and the results depicting the identified puncta were quantified using the *Analyze Particles* tool in Fiji.

### Immunoblots FLS2 and MPK

Seedlings grown on 0.5xMS agar with 0.5% sucrose for 9 days in continuous light (20°C) were transferred to 0.5xMS medium in 35 mm x 10 mm dishes so that there were 10-15 seedlings *per* genotype *per* treatment in each dish. These dishes were returned to continuous light on a shaker at 120 rpm overnight. On day 10, seedlings were treated +/− 100 nM flg22 (mock = PBS) +/− hypoxia in anaerojars (normoxia = ambient O_2_ levels) for 30 min or 1 hr for MAPK-phosphorylation and FLS2-GFP immunoblots, respectively. Seedlings were collected on liquid nitrogen and protein was extracted by grinding tissue in liquid nitrogen and adding 2xSDS loading buffer (Laemmli buffer) to the ground tissue in a 1:1 volume (μL):weight (mg) ratio. Proteins were denatured by heating samples to 95°C for 5 min followed by centrifuging at maximum speed for 10 min. The protein lysate was collected and protein concentration was determined using the amido black method (Schaffner and Weissmann, 1973; Popov et al., 1975). 30 µg of protein was loaded per well on 12% SDS-PAGE gels for MAPK-phosphorylation immunoblots; 100 µg of protein was loaded per well on 10% SDS-PAGE gels for FLS2-GFP immunoblots. Separated proteins were transferred from gels to PVDF membrane. Equal protein loading was assessed through Ponceau S staining (0.4% (w/v) Ponceau S, 10% (v/v) acetic acid in water)of the PVDF membrane. Membrane was blocked with 5% milk in TBS-T or PBS-T (containing 0.05% (v/v) Tween-20) for MAPK-phosphorylation and FLS2-GFP immunoblots, respectively. Blots were incubated overnight at 4°C with appropriate primary antibody with constant rotation: anti-MAPK-phosphorylation (Cell Signalling #4370S; 1:2,000); anti-GFP (Merck #1181446001; 1:1,000). Blots were washed with TBS-T or PBS-T for 3 x 5 minutes for MAPK-phosphorylation and FLS2-GFP blots, respectively, before and after incubation of blots with appropriate secondary antibody (anti-rabbit HRP (Merck #A0545; 1:50,000); anti-mouse HRP (Merck #A9044; 1:10,000)) for 2 hrs at room temperature. WesternBright ECL substrate (Advansta) was used and immunoblots were imaged using the G:BOX gel documentation system and the GeneSys software. Signal intensity of proteins of interest was quantified using Image J.

### Detection of reactive oxygen species

Flg22-induced reactive oxygen species (ROS) were detected using a luminol-based approach. Seeds were sown on 0.5xMS agar plates containing 0.5% sucrose, and stratified at 4°C for 3 days. Plates were transferred to a growth chamber at 20°C with 9 hrs of light *per* day. After 7 days of growth, individual seedlings were carefully moved to expanded Jiffy-7 (44 mm) pellets and grown in the same conditions for a further 3 weeks. Discs were taken from leaves of 4-week-old plants with a cork borer (1 cm diameter). Leaf discs were then carefully divided in 4 equal-sized quarters with a razor blade. Each quarter-disc was placed into a separate well of a white Sterilin 96-well plate (ThermoScientific) containing 200 μL dH_2_O with the abaxial leaf surface facing upwards. The plate was then returned to the growth room for a recovery period of at least 3 hours. 100x stock solutions of luminol (Merck) (17.7 mg/mL in 200 mM KOH) and horseradish peroxidase (HRP) (Fisher Scientific) (10 mg/mL in dH_2_O) were prepared fresh. 60 µL of a luminescence solution containing 2.8 µL 100X luminol, 2.8 µL 100X HRP and 54.4 µL dH_2_O was added to each well using a multichannel pipette. The plate was then transferred to a POLARstar Omega microplate reader (BMG LABTECH) and luminescence was detected for 15 minutes to establish a baseline measurement. After this time, 20 µL of flg22 solution (initial concentration: 1.4 μM; final concentration: 100 mM) was added to each well. Luminescence was detected every 2 minutes after addition of flg22.

## Acknowledgements

We thank Prof. Silke Robatzek for sharing the *FLS2_pro_:FLS2-3xmyc-GFP* line. This work was funded by grant 20/FFP-P/8433 from Science Foundation Ireland to EG. BCM was supported by an Irish Research Council PhD scholarship (GOIPG/2017/2); CMD was supported by an Irish Research Council PhD scholarship (GOIPG/2022/878) and by a PhD scholarship from the Kathleen Lonsdale Institute for Human Health Research (Maynooth University). Work in PFH lab was funded by the Deutsche Forschungsgemeinschaft (DFG, German Research Foundation, SFB1403 – Project-ID 414786233) and under Germany’s Excellence Strategy (CIBSS – EXC-2189 – Project ID 390939984). The authors confirm that they have no conflict of interest.

## Authors contributions

BCM, CMD, MM, PG, PFH and EG designed the work, conducted experiments, analyzed data and wrote the manuscript.

## Data Sharing

All data from this study are included in the article and/or in the supporting information. All lines are also available upon request. RNA-seq data have been deposited with the Gene Expression Omnibus (GEO) repository (at http://www.ncbi.nlm.nih.gov/) under GSE246173 and GSE246848. *Note to reviewers*: the data has been submitted and the authors are waiting for publication before release. Reviewer access can be requested. *Note to reviewers*: the RNA-seq data has been submitted and the authors are waiting for publication before public release. Reviewer access can be requested). The mass spectrometry proteomics data have been deposited to the ProteomeXchange Consortium (Deutsch et al., 2023) via the PRIDE (Perez-Riverol et al., 2022) partner repository with the dataset identifier PXD046263.

