## Supplementary Data for "Repression of pattern-triggered immune responses by hypoxia in Arabidopsis"

**Supplementary tables**

**Table S1: list of Arabidopsis accessions and genotypes used.**

| **Line** | **Description** | **Reference** |
| --- | --- | --- |
| Col-0 | Wild-type Columbia-0 accession |  |
| Ws | Wild-type Wassilewskija accession |  |
| *ate1-2 ate2-1* | SALK_023492 x SALK_040788 | (Graciet et al., 2009) |
| *prt6-1* | SAIL_1278_H11 | (Garzon et al., 2007) |
| *prt6-5* | SALK_051088 | (Graciet et al., 2009) |
| *ATE1 rescue in ate1ate2* | *ATE1* genomic locus restored in *ate1 ate2* background | (Graciet et al., 2009) |
| *erfVII* | Quintuple mutant for *RAP2.2/3/12* and *HRE1/2* | (Abbas et al., 2015) |
| *prt6-1 erf VII* | Sextuple mutant | (Abbas et al., 2015) |

**Table S2: Oligonucleotides used in this study for qPCR.**

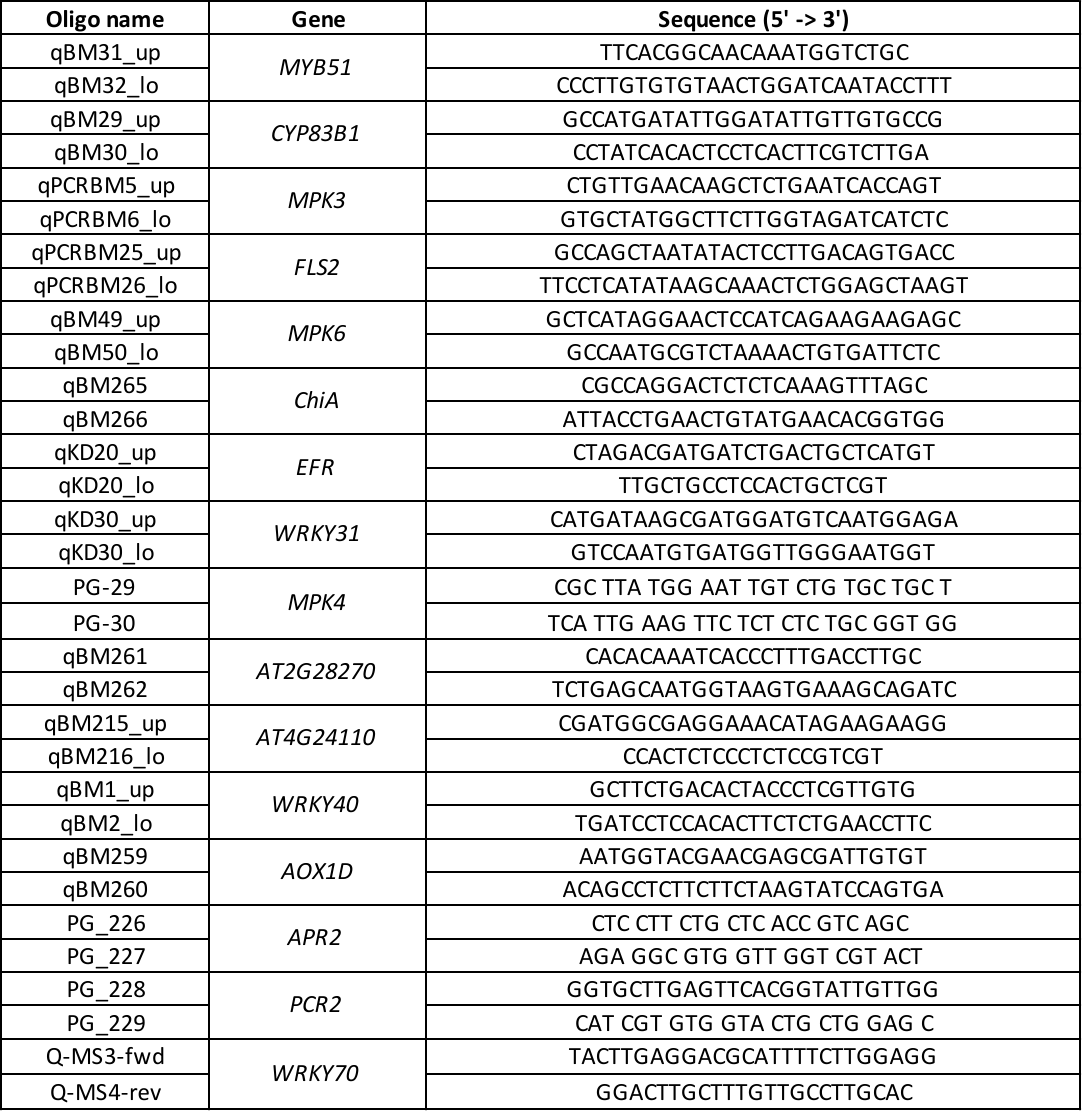

**Supplemental Figures**

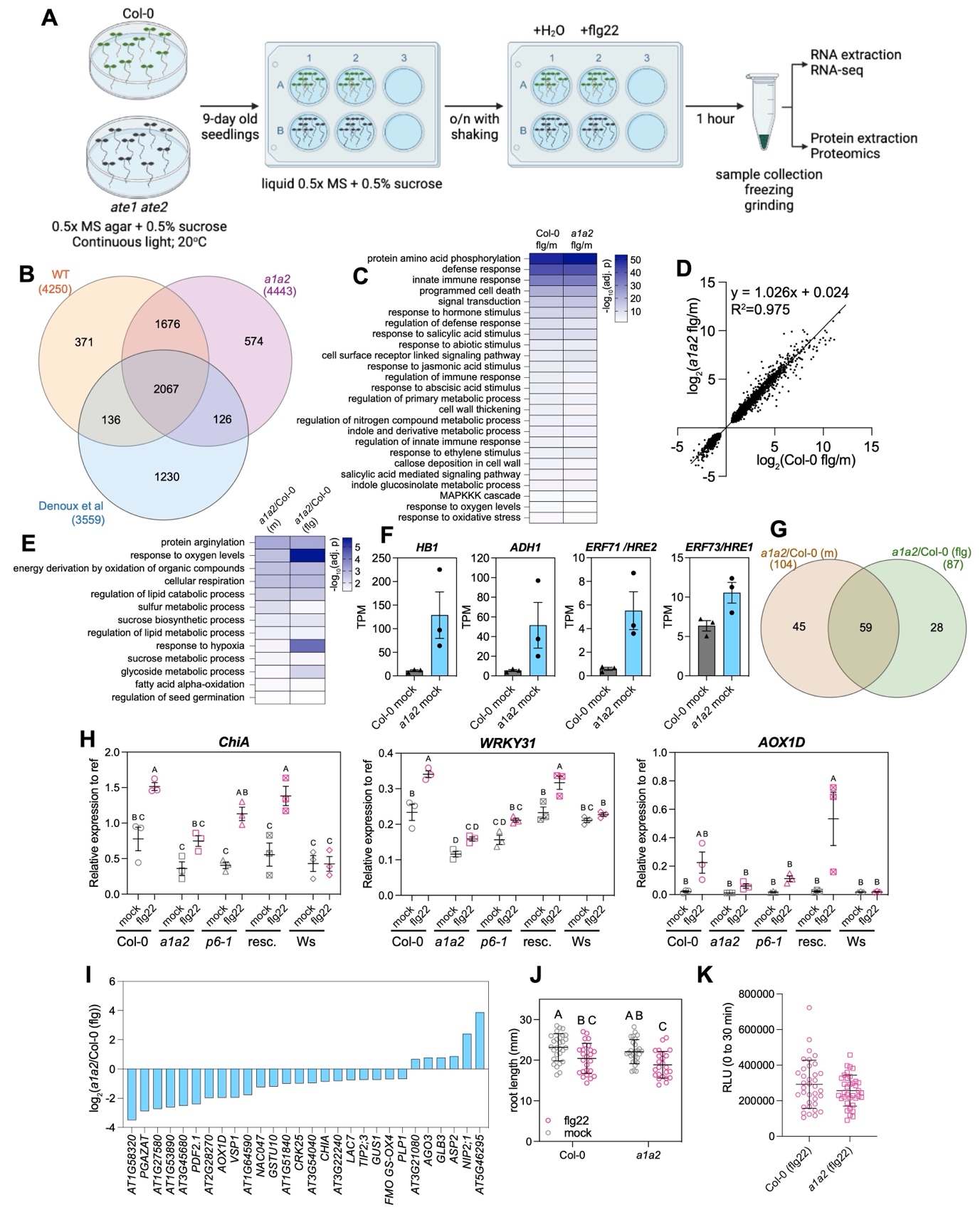

**Figure S1: Analyses of RNA-seq datasets with Col-0 and *ate1 ate2* mutants.**

**(A)** Overview of the experimental design to compare the transcriptomes and proteomes of wild-type Col-0 and *a1a2* mutant seedlings treated with a mock solution or with 1 µM flg22. RNA and proteins were extracted from the same sample for transcriptomic and proteomic analyses. **(B)** Overlap analysis between the flg/m datasets in wild-type (Col-0) and *a1a2* seedlings, and a previously published dataset (Denoux et al.) generated using similar experimental conditions (Col-0 accession; 1 µM flg22 for 1 hr) (Denoux et al., 2008). Cut-offs: adj. *p*-value<0.05 and |log2(FC)|>0.585). **(C)** Selected GO terms enriched in both the wild-type and *a1a2* mutant datasets, when comparing flg22 and mock-treated samples. See Suppl. Dataset 1 for the full list of GO terms. **(D)** Comparison of the amplitude of gene expression change between *a1a2* flg/m and Col-0 flg/m for DEGs common to both datasets. **(E)** Selected GO categories over-represented in the *a1a2*/Col-0 (m) and *a1a2*/Col-0 (flg) datasets. See Suppl. Dataset 1 for the full list of GO terms. **(F)** Expression of selected hypoxia-response genes in the *a1a2*/Col-0 (mock) RNA-seq dataset. Mean TPM (transcript per million) values and SEM of 3 biological replicates are shown. **(G)** Overlap analysis between *a1a2*/Col-0 (m) and *a1a2*/Col-0 (flg). **(H)** RT-qPCR analysis of the expression of selected flg22 responsive genes in wild-type Col-0, *a1a2*, *prt6-1* (*p6-1*), an *a1a2* rescue line ((Graciet et al., 2009); labelled ‘resc.’) that contains a transgene to express *ATE1* from its endogenous genomic locus, and Ws which lacks *FLS2* (Zipfel et al., 2004). Mean and SEM of 3 biological replicates are shown, including the results of two-way ANOVA and Tukey’s test. **(I)** Differential expression of the 28 unique DEGs in *a1a2* compared to the wild type in the presence of flg22. **(J)** Root growth inhibition assays. Root elongation on vertical plates was measured with Image J during a 3-day period following the addition of 500 nM flg22. Means and standard deviations are shown from 4 biological replicates with 4 to 7 seedlings *per* genotype *per* treatment *per* replicate. The results of two-way ANOVA with Tukey’s multiple comparison are shown. **(K)** Apoplastic ROS production in response to flg22. Sum of luminescence over a 30-min period after the addition of flg22 is shown. Five biological replicates were carried out with 6-8 leaf discs for each genotype and in each biological replicate. Means and standard deviations are shown. No statistically significant difference.

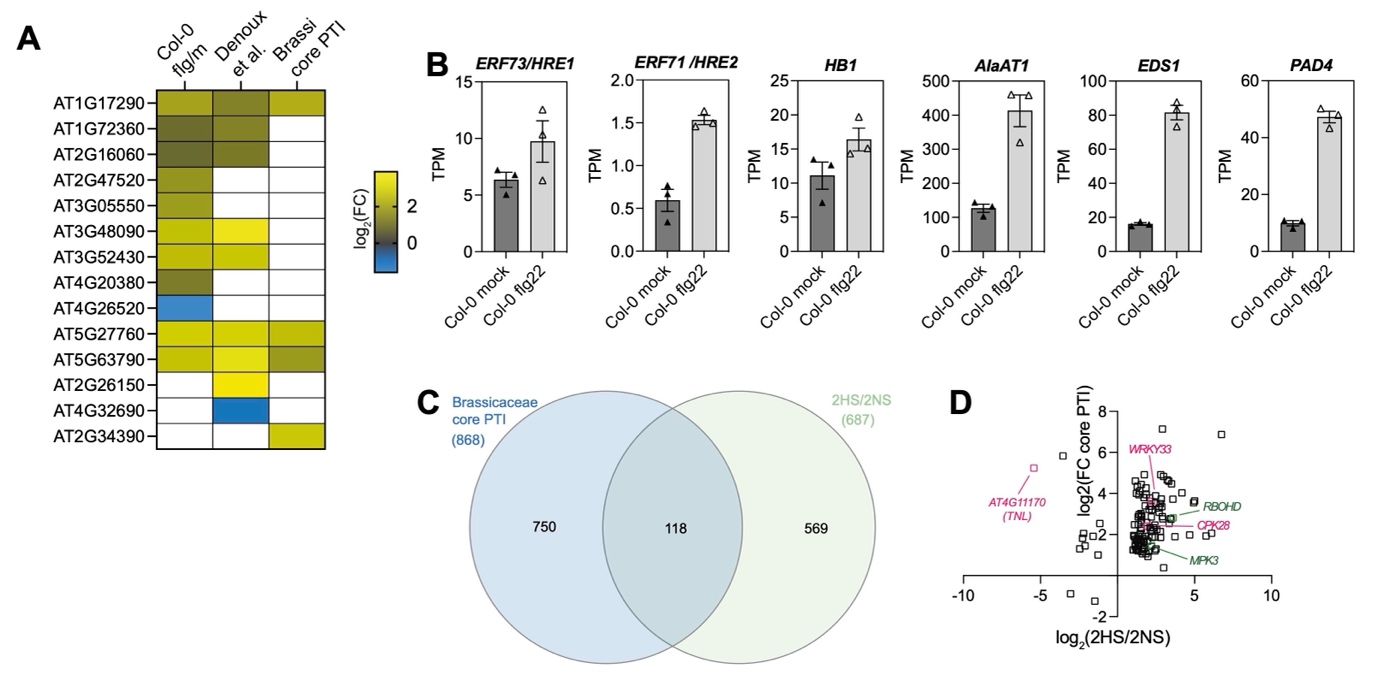

**Figure S2: Overlap between flg22 and hypoxia response programs.**

**(A)** Gene expression changes shown as log_2_(fold change) for ‘response to oxygen levels’ and ‘response to hypoxia’ DEGs in the different datasets. **(B)** Expression of selected hypoxia-response genes in the Col-0 flg/m RNA-seq dataset. Mean TPM (transcript per million) values and SEM of 3 biological replicates are shown. **(C)** Overlap between Brassicaceae core PTI genes (Winkelmuller et al., 2021) and hypoxia response genes (2HS/2NS (Lee and Bailey-Serres, 2019)). **(D)** Comparison of the directionality and amplitude of gene expression change for the 118 genes common to Brassicaceae core PTI and 2HS/2NS.

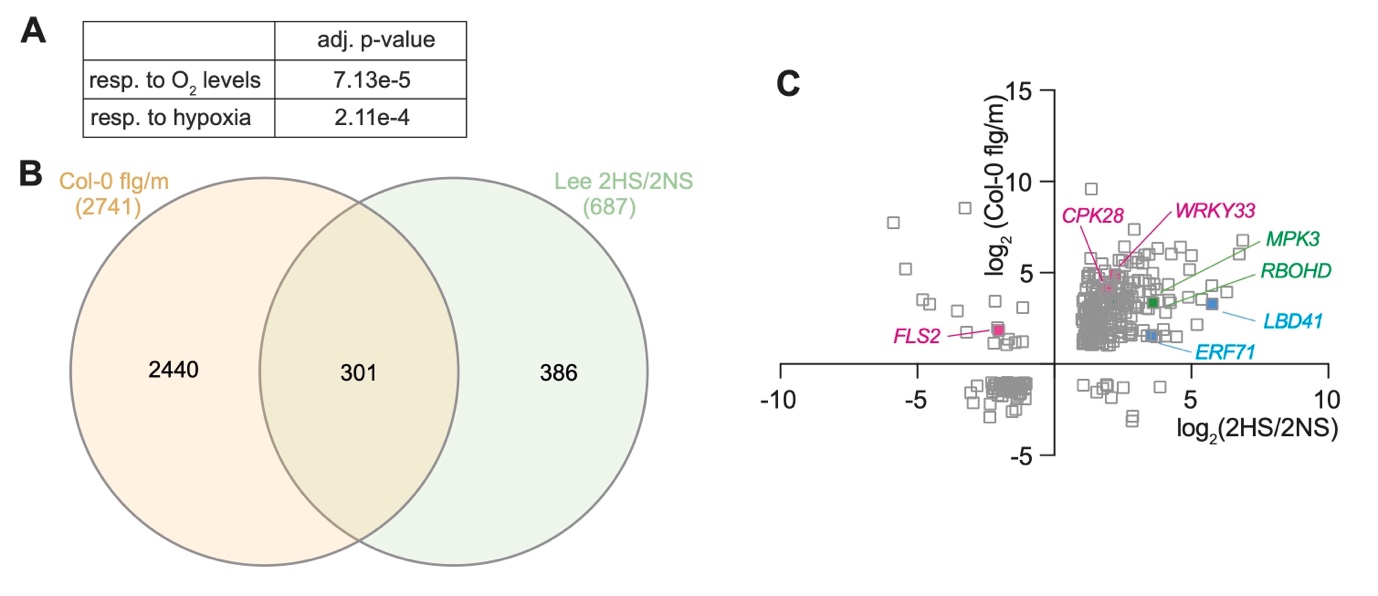

**Figure S3:** **Overlap between the flg22 and hypoxia transcriptional response programs with more stringent cut-offs on Col-0 flg/m.** Here, |log_2_(FC)|>1.0 and adj. *p*-value<0.05 were applied to the Col-0 flg/m dataset. **(A)** Enrichment for GO categories associated with oxygen levels: ‘response to oxygen levels’ (GO#70482) and ‘response to hypoxia’ (GO#1666). **(B)** Overlap between flg22 response genes identified in Col-0 and differentially regulated genes after 2 hrs of hypoxia (2HS/2NS; O_2_<2%; cut-off applied: |log_2_(FC)|>1.0 and FDR<0.05) (Lee and Bailey-Serres, 2019). **(C)** Comparison of the directionality and amplitude of gene expression change for the 301 genes common to Col-0 flg/m and 2HS/2NS.

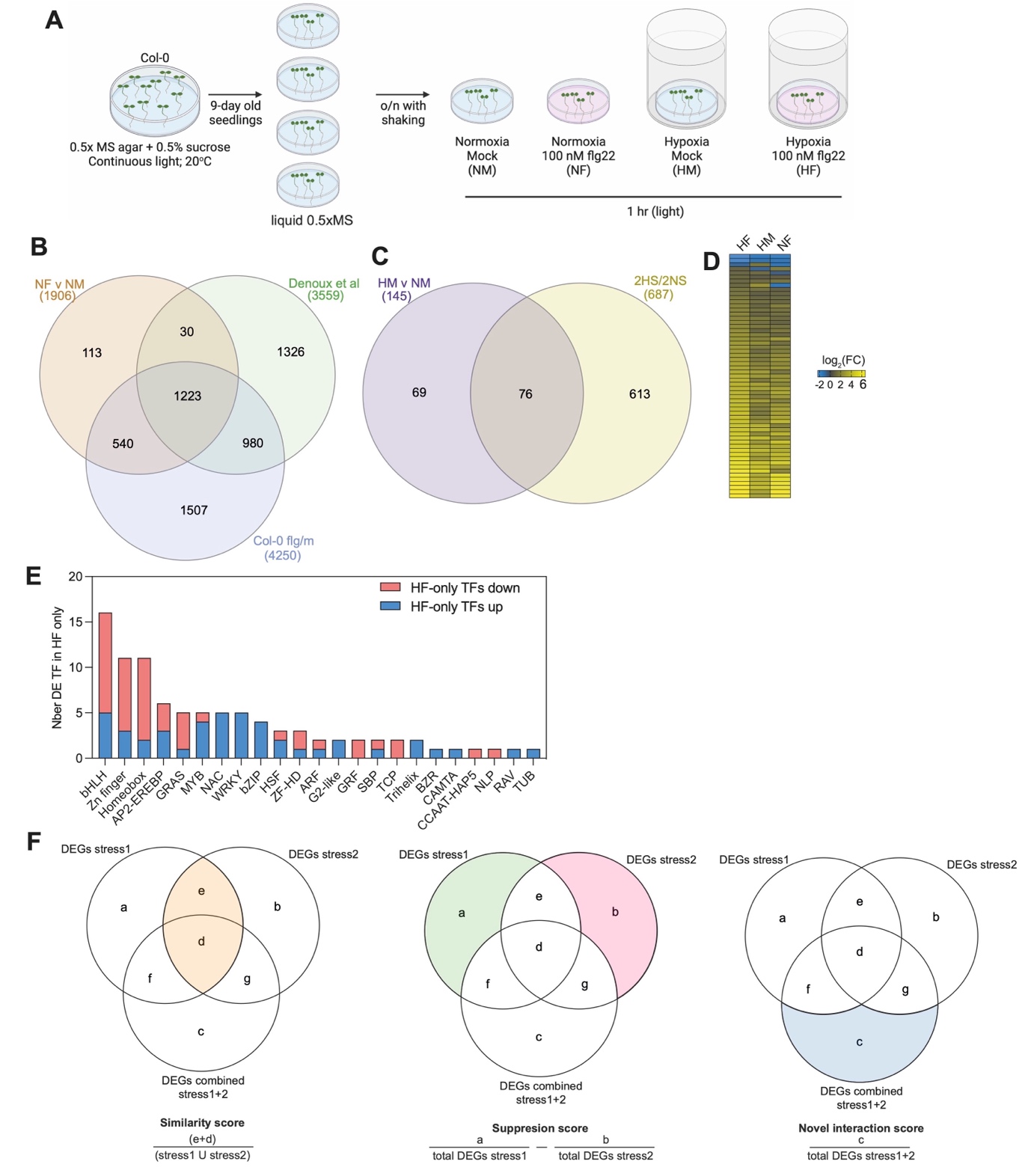

**Figure S4: Analysis of transcriptional changes under combined hypoxia/flg22.**

**(A)** Experimental design of the RNA-seq experiment to compare gene expression changes between individual hypoxia or flg22 treatments and combined hypoxia/flg22 treatment. **(B)** Flg22-response genes (NF v NM; 100 nM flg22 ) were compared to those obtained in response to flg22 in (Denoux et al., 2008) (1 µM flg22) and in our Col-0 flg/m (1 µM flg22). **(C)** Hypoxia response DEGs (HM v NM) were compared to those obtained in (Lee and Bailey-Serres, 2019) after 2 hrs of hypoxia (2HS/2NS). **(D)** Comparison of the gene expression changes for the 59 DEGs common to all 3 datasets. **(E)** Transcription factor families and corresponding number of up- or down-regulated transcription factors among the 939 DEGs unique to combined hypoxia/flg22 treatment. **(F)** Calculation of metrics to characterize the transcriptional response to combined stresses, as outlined in (Tan et al., 2023). Single letters in the different sections of the Venn diagrams correspond to the number of DEGs in that particular section. Stress1 U stress2: total number of DEGs in stress1 and stress2.

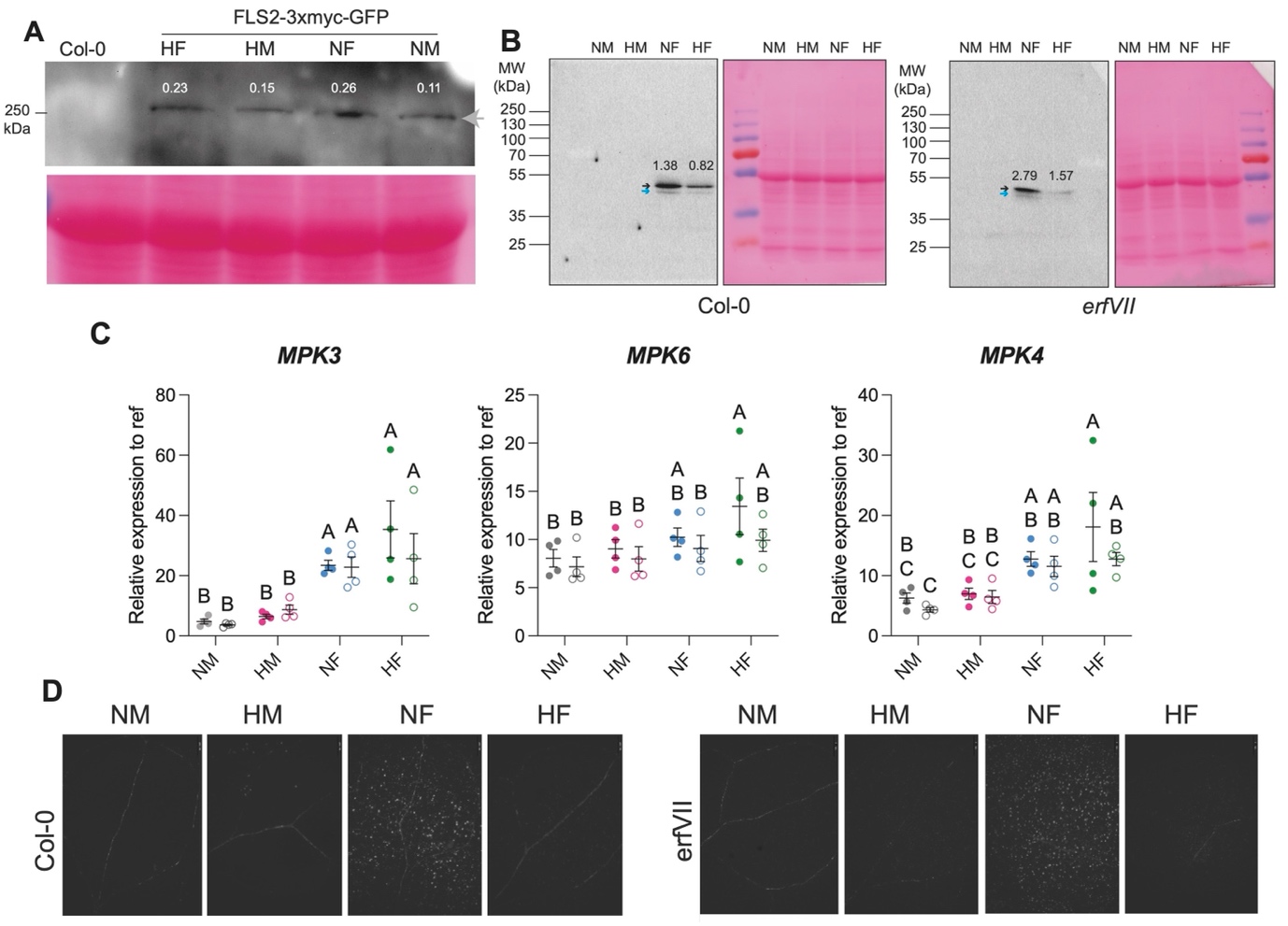

**Figure S5: Effects of combined hypoxia/flg22 on PTI.**

**(A)** Immunoblot analysis of FLS2 levels in a *FLS2pro:FLS2-3xmyc-GFP* line (Robatzek et al., 2006). The grey arrow indicates FLS2-3xmyc-GFP (MW: 164 kDa). Ponceau-normalized signal intensity determined using Image J for FLS2-3xmyc-GFP are indicated above the bands. **(B)** Immunoblots and corresponding Ponceau stainings for anti-phosphorylated MPK immunoblots shown in Fig. 4. Black arrow: MPK6 (45 kDa); blue arrow: MPK3 (43 kDa). Ponceau-normalized signal intensity determined using Image J for MPK3/6 together are indicated above the bands. **(C)** Relative expression of *MPK3/4/6*. Each data point, mean and standard deviations are shown from 4 biological replicates with 10 seedlings for each genotype and condition in one given biological replicate, with the exception of one biological replicate with 5 seedlings for all samples. The results of two-way ANOVA and Fisher’s tests are shown. **(D)** Representative pictures of callose deposition assays to compare the effects of hypoxia/flg22 to those of normoxia/flg22 in Col-0 and *erfVII*.
